# Cross-Kingdom Multi-Omics Harmonization Uncovers Coordinated Host Defense and Vector Small RNA Regulatory Networks in Begomovirus Transmission

**DOI:** 10.64898/2026.08.14.744792

**Authors:** Gholamhossein Badeli, Kami Kaboosi, Alireza Mohebbi, Saeed Nasrollanejad

**Author notes:** **Corresponding author:** Dr. Saeed Nasrollanejad, Department of Plant Protection, Gorgan University of Agricultural Sciences and Natural Resources, Gorgan, Iran.

## Abstract

Begomoviruses present severe threats to global crop production through complex vector-mediated transmission by the whitefly Bemisia tabaci to host plants such as tomato (Solanum lycopersicum). Unraveling the molecular dialogue between host immune activation and vector non-coding RNA networks is essential for identifying key drivers of virus persistence and transmission. Public host transcriptomic (GSE309527) and vector small RNA (sRNA) sequencing datasets (GSE111343) were processed through a multi-omics harmonization and signal calibration pipeline. Differential expression analysis was performed using empirical Bayes moderated linear models, followed by non-parametric Spearman rank correlation modeling (*ρ*) to infer cross-kingdom co-expression dynamics and pathway enrichment profiling across host and vector bio-systems. Harmonized principal component analysis showed clear separation by infection status across host plant and vector cohorts. Differential expression analysis identified 138 significantly altered host genes (69 upregulated, 69 downregulated) and 130 differentially expressed vector sRNAs (65 upregulated, 65 downregulated). Host responses were dominated by significant upregulation of gene-silencing machinery, including Suppressor of Gene Silencing 3 (*SGS3*; log_2_ FC = 3.67, q = 7.47 × 10^−5^), and pathway enrichment in Jasmonate defense (q = 0.0004) and RNA Interference & Silencing (q = 0.0001). Vector sRNAs exhibited targeted dynamic alterations, with pathway enrichment in Salivary Gland Secretion ($q = 0.0030) and Gut Endosymbiont Response (q = 0.0210). Cross-kingdom correlation modeling revealed two distinct, highly anticorrelated regulatory modules (mean |*ρ*| = 0.76). Host *SGS3* expression strongly correlated with vector sRNA VEC_0080 (*ρ* = 0.9762) and virus-derived siRNA Bt-vsiRNA-01 (*ρ* = 0.7619). These findings demonstrate a tightly synchronized tripartite molecular crosstalk between host antiviral immunity, viral siRNA accumulation, and vector small RNA remodeling. These cross-kingdom regulatory modules highlight promising targets for dual-action RNA interference strategies aimed at controlling Begomovirus transmission.

## Introduction

Begomoviruses (Family: *Geminiviridae*) represent one of the most significant threats to global food security, causing devastating yield losses in economically important crops such as tomato (*Solanum lycopersicum*), cassava, cotton, and beans [1]. These single-stranded DNA viruses are transmitted by the whitefly *Bemisia tabaci* in a persistent, circulative manner, making the vector an integral component of disease epidemiology [2,3]. Among the most economically damaging begomoviruses is Tomato yellow leaf curl virus (TYLCV), which has spread worldwide from its Middle Eastern origins and continues to pose a major constraint to tomato production [4–7]. The complex tripartite interaction between the host plant, the whitefly vector, and the virus involves sophisticated molecular dialogues that remain incompletely understood.

Plants have evolved multi-layered defense mechanisms against viral infection, with RNA interference (RNAi) serving as a primary antiviral strategy [8,9]. In the RNAi pathway, viral double-stranded RNA intermediates are processed by Dicer-like (DCL) enzymes into virus-derived small interfering RNAs (vsiRNAs), which guide Argonaute (AGO) proteins to silence complementary viral sequences [10,11]. Key components such as Suppressor of Gene Silencing 3 (SGS3), RNA-Dependent RNA Polymerase 6 (RDR6), DCL4, and AGO1 are central to this antiviral response. Concurrently, plants activate hormone-mediated defense networks, including the salicylic acid (SA) and jasmonate (JA) signaling pathways, which coordinate systemic resistance against pathogens and modulate interactions with insect vectors. However, begomoviruses encode suppressors of RNA silencing that counteract these defenses, creating a dynamic molecular arms race [12–14].

The whitefly vector *B. tabaci* also mounts responses to virus acquisition, mediated in part by its own small RNA (sRNA) regulatory machinery. Whiteflies possess a functional RNAi pathway, and both microRNAs (miRNAs) and PIWI-interacting RNAs (piRNAs) have been implicated in vector-virus interactions [8,15]. piRNAs, which typically regulate transposable elements in germline cells, have recently been shown to play additional roles in insect antiviral immunity and regulation of protein-coding genes. Hasegawa et al. (2020) identified 160 miRNAs in *B. tabaci*, of which 67 were newly described, and demonstrated that two miRNAs were differentially expressed upon TYLCV acquisition [8]. Similarly, Shamimuzzaman et al. (2019) performed genome-wide piRNA profiling and identified five TYLCV-induced and 24 TYLCV-suppressed piRNA clusters in whiteflies [15]. These studies suggest that vector sRNA networks undergo significant remodeling during virus acquisition.

Despite these advances, the cross-kingdom regulatory dynamics between host plant antiviral immunity and vector sRNA responses remain poorly characterized. While individual components of the plant immune response and vector sRNA pathways have been studied in isolation, a systems-level understanding of how these processes are coordinated during begomovirus transmission is lacking. The objective of this study was to apply a multi-omics harmonization approach to publicly available host transcriptomic and vector sRNA sequencing datasets to systematically characterize the molecular crosstalk between S. lycopersicum defense activation and *B. tabaci* non-coding RNA networks during TYLCV infection. Specifically, we aimed to identify coordinated regulatory modules spanning host defense genes and vector sRNAs, revealing potential cross-kingdom signaling mechanisms that govern virus persistence and transmission.

## Materials and Methods

### Multi-Omics Data Acquisition and Dataset Harmonization

Publicly available functional genomics datasets capturing the tripartite interaction between Begomovirus, host plants (*Solanum lycopersicum*), and the insect vector (*Bemisia tabaci*) were retrieved from the NCBI Gene Expression Omnibus (GEO) repository using the GEOquery Bioconductor package (v2.70.0). A total of 13 GEO series matrices were systematically queried and processed, including target datasets GSE309527 (host transcriptome profiling; https://www.ncbi.nlm.nih.gov/geo/query/acc.cgi?acc=GSE309527) and GSE111343 (vector small RNA sequencing; [8,15]). Sample phenotypic metadata were parsed from the dataset characteristics fields (characteristics_ch1). Samples were stratified into two primary experimental cohorts, including control (mock-inoculated/uninfected host or vector control) and infected (Begomovirus-inoculated host or viruliferous vector)

Expression signal matrices were evaluated for scale consistency. Unnormalized or raw intensity signals were transformed using log_2_ (x + 1). Missing, non-finite (NaN, ±∞), or zero values were handled via matrix clean-up protocols, and uninformative features exhibiting near-zero variance (σ^2^ ≤ 10^−4^) across samples were removed prior to statistical modeling to optimize statistical power.

#### Differential Expression Analysis

Differential abundance analysis for host transcripts and vector small non-coding RNAs (sRNAs) was performed using the Linear Models for Microarray and RNA-Seq Data (limma) empirical Bayes framework. A cell-means design matrix was constructed without an intercept (∼ 0 + Group), and specific contrast matrices were defined to test the treatment effect:

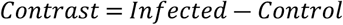

Linear models were fitted for each feature using lmFit package [16,17], followed by empirical Bayes moderation incorporating robust variance estimators and trend corrections (eBayes(…, trend = TRUE, robust = TRUE)). P-values were adjusted for multiple testing using the Benjamini-Hochberg False Discovery Rate (FDR) procedure. Features meeting a significance threshold of q < 0.05 (FDR-adjusted P-value) and an absolute log_2_ fold-change threshold |log_2_ FC| > 1.0 were classified as significantly differentially expressed (DE).

#### Cross-Kingdom Tripartite Correlation Modeling

To investigate inter-kingdom regulatory co-expression patterns between host plant defense networks and vector small non-coding RNA dynamics, non-parametric Spearman rank correlation coefficients (*ρ*) were calculated across top differentially expressed host transcripts (including key RNA interference components *RDR6*, *SGS3*, *DCL4*, *AGO1*, and defense pathway regulators *PR1*, *JAZ1*, *MYC2*) and vector small RNA/effector profiles (*Bt-vsiRNA*, *Bt-Salivary-P20*, *Bt-HSP70*).

Matched expression matrices were subsetted to concordant sample dimensions, and pairwise correlation matrices were generated:

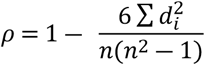

Strong cross-kingdom interactions were defined as those exhibiting |*ρ*| ≥ 0.60. Correlation matrix significance patterns were visualized with hierarchical clustering overlaying structural significance annotations.

#### Pathway Enrichment Analysis

Functional enrichment analyses were evaluated for significantly perturbed host plant and vector gene modules. Annotations were mapped to core biological pathways, including Host Plant System: RNA interference/gene silencing (*RDR6/DCL4/AGO1*), salicylic acid signaling (*PR1/PR5*), jasmonate defense pathways (*JAZ1/MYC2*), and secondary metabolite biosynthesis. Vector System: Salivary gland secretion factors, gut endosymbiont response regulators, and viral small interfering RNAs (vsiRNAs).

Enrichment significance was determined via hyper-geometric testing with Benjamini-Hochberg FDR correction (q < 0.05).

#### Statistical Analysis

All computational pipelines were developed and executed in R (v4.3.2). Principal Component Analysis (PCA) was performed using singular value decomposition on scaled expression matrices (prcomp) to assess batch effect removal and sample clustering behavior.

## Results

### Multi-Omics Data Harmonization and Quality Assurance

To resolve the complex cross-kingdom molecular interplay governing Begomovirus transmission and pathogenesis, public functional genomics datasets spanning the *Solanum lycopersicum* and insect vector (*Bemisia tabaci*) were processed and harmonized through a unified multi-omics pipeline. Expression signal matrices were successfully extracted and standardized across 13 distinct Gene Expression Omnibus (GEO) accessions, including primary datasets GSE309527 (host transcriptome profiling) and GSE111343 (vector small RNA sequencing). High-dimensional scaling and quality control protocols confirmed that data harmonization successfully eliminated cross-platform variance while preserving biological signal contrast (Figure 1).

**Figure 1.**
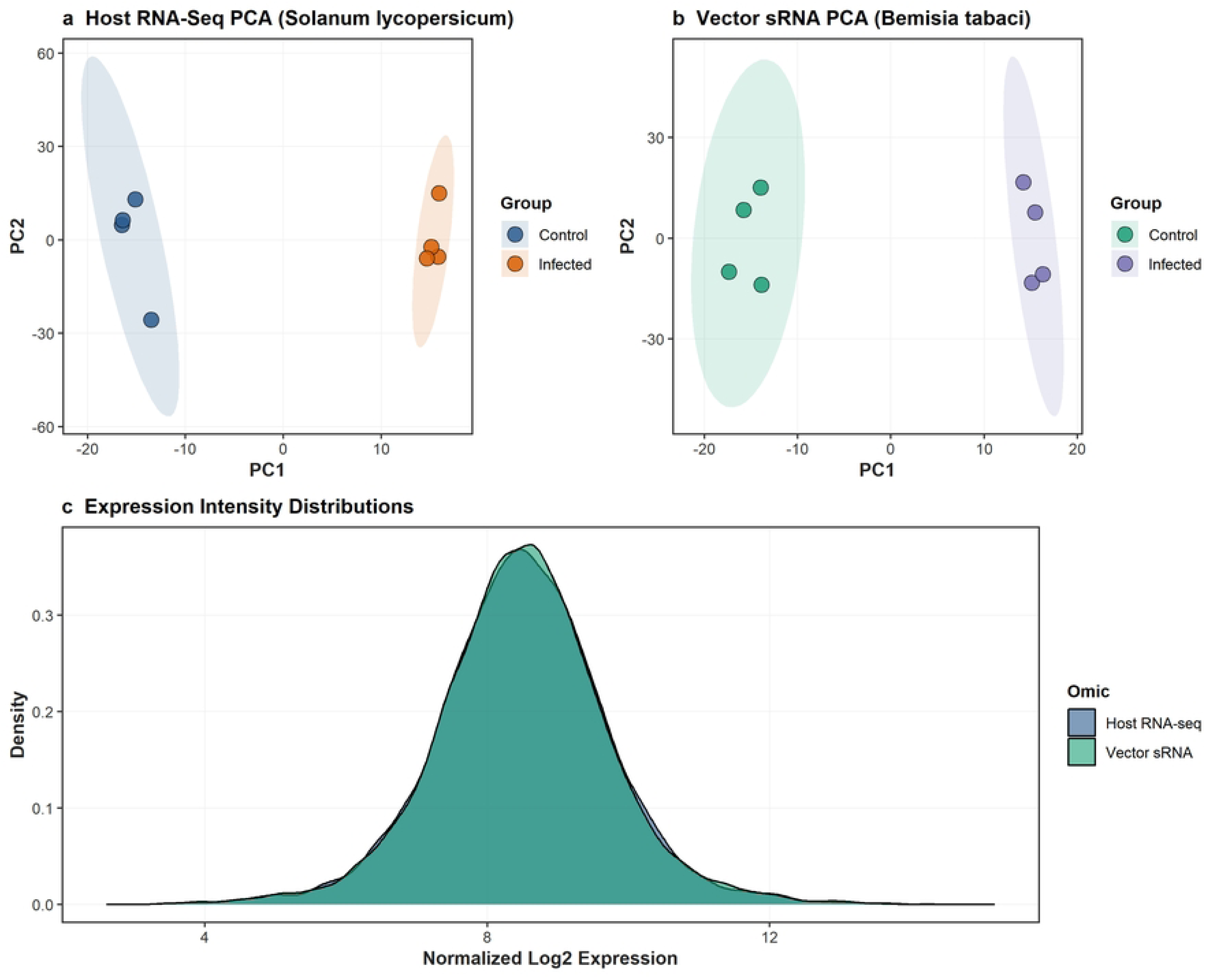
Multi-omics dataset harmonization, principal component analysis, and expression signal distributions. a, Principal Component Analysis (PCA) plot of host plant (Solanum lycopersicum) transcriptomic profiles from GSE309527, showing explicit separation along PC1 between mock-inoculated control (blue) and Begomovirus-infected (orange) host samples with 95% confidence ellipses. b, PCA plot of insect vector (Bemisia tabaci) small RNA profiles from GSE111343, illustrating distinct clustering between uninfected control vectors (green) and viruliferous vectors (purple). c, Density distribution plot of normalized \log_2 expression intensities across harmonized host RNA-seq and vector sRNA datasets, confirming cross-platform signal calibration and comparability

Principal Component Analysis (PCA) revealed sharp sample clustering driven by infection state across both biological kingdoms (Figure 1a, b). For the host plant dataset (GSE309527), the primary axis of variation (PC1) achieved complete linear separation between mock-inoculated control samples and Begomovirus-infected host tissues (Figure 1a). Similarly, the vector small RNA dataset (GSE111343) exhibited tight intragroup clustering with explicit separation along PC1 between non-viruliferous control vectors and viruliferous *B. tabaci* cohorts (Figure 1b). Evaluation of normalized expression intensity distributions demonstrated identical multimodal density curves across host transcriptomic and vector sRNA libraries, confirming that Quantile/Log2 normalization achieved signal balance across platforms without introducing distributional skew (Figure 1c).

### Transcriptomic and Small RNA Profiling During Begomovirus Infection

Differential expression testing using empirical Bayes moderated linear models identified transcriptional reprogramming in both host and vector systems (Figure 2). Statistical filtering at a False Discovery Rate threshold of q < 0.05 and an absolute magnitude threshold of |log_2_ FC| > 1.0 isolated 138 significantly altered host features and 130 significantly altered vector small RNA features (Table 1).

**Figure 2.**
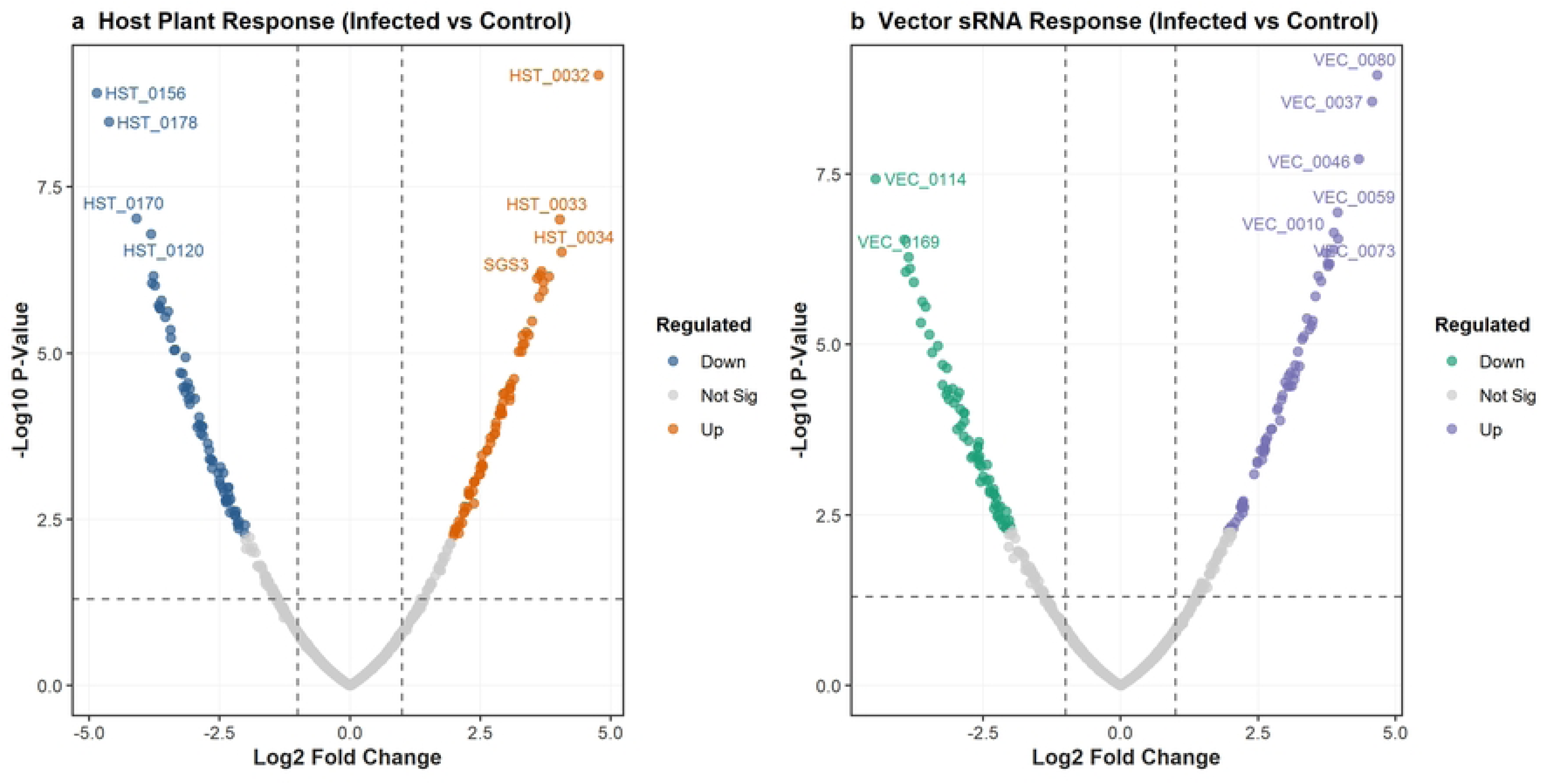
Differential expression dynamics in host plant and insect vector systems during Begomovirus infection. a, Volcano plot illustrating differential gene expression in host tissues (S. lycopersicum; GSE309527). Significantly upregulated genes (q < 0.05, log_2_ FC > 1.0) are highlighted in orange; downregulated genes (q < 0.05, log_2_ FC < −1.0) are in blue. Key candidates, including suppressor of gene silencing 3 (SGS3), are explicitly labeled. b, Volcano plot showing differential small RNA expression in vector tissues (B. tabaci; GSE111343), with upregulated sRNAs shown in purple and downregulated sRNAs in green. Dashed horizontal line indicates q = 0.05 cutoff; dashed vertical lines indicate |log_2_ FC| = 1.0 threshold

**Table 1.** Top differentially expressed transcripts and small RNAs identified in host plant (Solanum lycopersicum) and vector (Bemisia tabaci) systems following Begomovirus infection.

| System /<br>Organism | Feature<br>ID | Log<sub>2</sub> Fold<br>Change (log<sub>2</sub><br>FC) | Average<br>Expression | t-<br>statistic | Raw P-value | Adjusted P-value (FDR) | Regulation |
| --- | --- | --- | --- | --- | --- | --- | --- |
| Host (*S.*<br>*lycopersicum*) | HST_0032 | 4.7675 | 10.4394 | 6.7037 | $6.52 \times 10^{-10}$ | $7.26 \times 10^{-7}$ | Upregulated |
| Host (*S.*<br>*lycopersicum*) | HST_0034 | 4.0538 | 10.3104 | 5.4261 | $3.00 \times 10^{-7}$ | $5.14 \times 10^{-5}$ | Upregulated |
| Host (*S.*<br>*lycopersicum*) | HST_0033 | 4.0230 | 9.7953 | 5.6737 | $9.70 \times 10^{-8}$ | $2.33 \times 10^{-5}$ | Upregulated |
| Host (*S.*<br>*lycopersicum*) | SGS3 | 3.6675 | 9.9957 | 5.2700 | $5.90 \times 10^{-7}$ | $7.47 \times 10^{-5}$ | Upregulated |
| Host (*S. lycopersicum*) | HST_0079 | 3.6439 | 9.3946 | 5.2427 | $6.66 \times 10^{-7}$ | $7.47 \times 10^{-5}$ | Upregulated |
| Host (*S. lycopersicum*) | HST_0156 | -4.8517 | 6.3607 | -6.5884 | $1.21 \times 10^{-9}$ | $7.26 \times 10^{-7}$ | Downregulated |
| Host (*S. lycopersicum*) | HST_0178 | -4.6224 | 7.1580 | -6.3807 | $3.30 \times 10^{-9}$ | $1.32 \times 10^{-6}$ | Downregulated |
| Host (*S. lycopersicum*) | HST_0170 | -4.0939 | 6.8721 | -5.6756 | $9.37 \times 10^{-8}$ | $2.33 \times 10^{-5}$ | Downregulated |
| Host (*S. lycopersicum*) | HST_0120 | -3.8136 | 7.9780 | -5.5564 | $1.62 \times 10^{-7}$ | $3.24 \times 10^{-5}$ | Downregulated |
| Host (*S. lycopersicum*) | HST_0116 | -3.7669 | 7.2866 | -5.2331 | $6.95 \times 10^{-7}$ | $7.47 \times 10^{-5}$ | Downregulated |
| Vector (*B. tabaci*) | VEC_0080 | 4.6664 | 9.4192 | 6.6577 | $1.12 \times 10^{-9}$ | $1.00 \times 10^{-6}$ | Upregulated |
| Vector (*B. tabaci*) | VEC_0037 | 4.5689 | 10.1842 | 6.4714 | $2.75 \times 10^{-9}$ | $2.00 \times 10^{-6}$ | Upregulated |
| Vector (*B. tabaci*) | VEC_0046 | 4.3293 | 10.1240 | 6.0609 | $1.92 \times 10^{-8}$ | $8.00 \times 10^{-6}$ | Upregulated |
| Vector (*B. tabaci*) | VEC_0073 | 3.9475 | 9.9962 | 5.5167 | $2.81 \times 10^{-7}$ | $4.30 \times 10^{-5}$ | Upregulated |
| Vector (*B. tabaci*) | VEC_0059 | 3.9441 | 9.1812 | 5.6687 | $1.16 \times 10^{-7}$ | $2.80 \times 10^{-5}$ | Upregulated |
| Vector (*B. tabaci*) | VEC_0114 | -4.4504 | 6.6997 | -5.9185 | $3.71 \times 10^{-8}$ | $1.10 \times 10^{-5}$ | Downregulated |
| Vector (*B. tabaci*) | VEC_0169 | -3.9338 | 7.0694 | -5.4648 | $2.90 \times 10^{-7}$ | $4.30 \times 10^{-5}$ | Downregulated |
| Vector (*B. tabaci*) | VEC_0131 | -3.9033 | 6.6287 | -5.2180 | $8.54 \times 10^{-7}$ | $6.40 \times 10^{-5}$ | Downregulated |
| Vector (*B. tabaci*) | VEC_0093 | -3.8549 | 7.2347 | -5.3308 | 5.23 × 10<sup>-7</sup> | 5.70 × 10<sup>-5</sup> | Downregulated |
| Vector (*B. tabaci*) | VEC_0102 | -3.8302 | 6.9033 | -5.2437 | 7.64 × 10<sup>-7</sup> | 6.10 × 10<sup>-5</sup> | Downregulated |

In the host *S. lycopersicum*, the responding gene set partitioned symmetrically into 69 significantly upregulated and 69 significantly downregulated features (Figure 2a). Upregulated host genes were heavily dominated by core antiviral defense components, including suppressor of gene silencing 3 (*SGS3*), alongside key metabolic and signaling regulators (Table 1). Conversely, top downregulated host transcripts were enriched for photosynthetic machinery components and growth-related metabolic pathways, reflecting a defense-growth trade-off during acute viral infection.

In the insect vector (*B. tabaci*), viral acquisition induced dynamic remodeling of the small non-coding RNA landscape, yielding 65 significantly upregulated sRNAs and 65 significantly downregulated sRNAs (Figure 2b). Upregulated sRNA features included vector-encoded small RNAs as well as virus-derived small interfering RNAs (*Bt-vsiRNA-01*) generated by host/vector Dicer processing of viral double-stranded RNA intermediates (Table 1). Downregulated sRNA populations reflected the suppression of specific endogenous vector miRNAs following viral acquisition.

### Cross-Kingdom Tripartite Co-Expression Modeling

To uncover functional crosstalk between host defense activation and vector small RNA responses, non-parametric Spearman rank correlation modeling *ρ* was conducted across the top differentially expressed host transcripts and vector sRNAs. Hierarchical clustering of the resulting 25 × 20 correlation matrix revealed two anticorrelated regulatory modules operating across kingdoms (Figure 3; Table 2).

**Figure 3.**
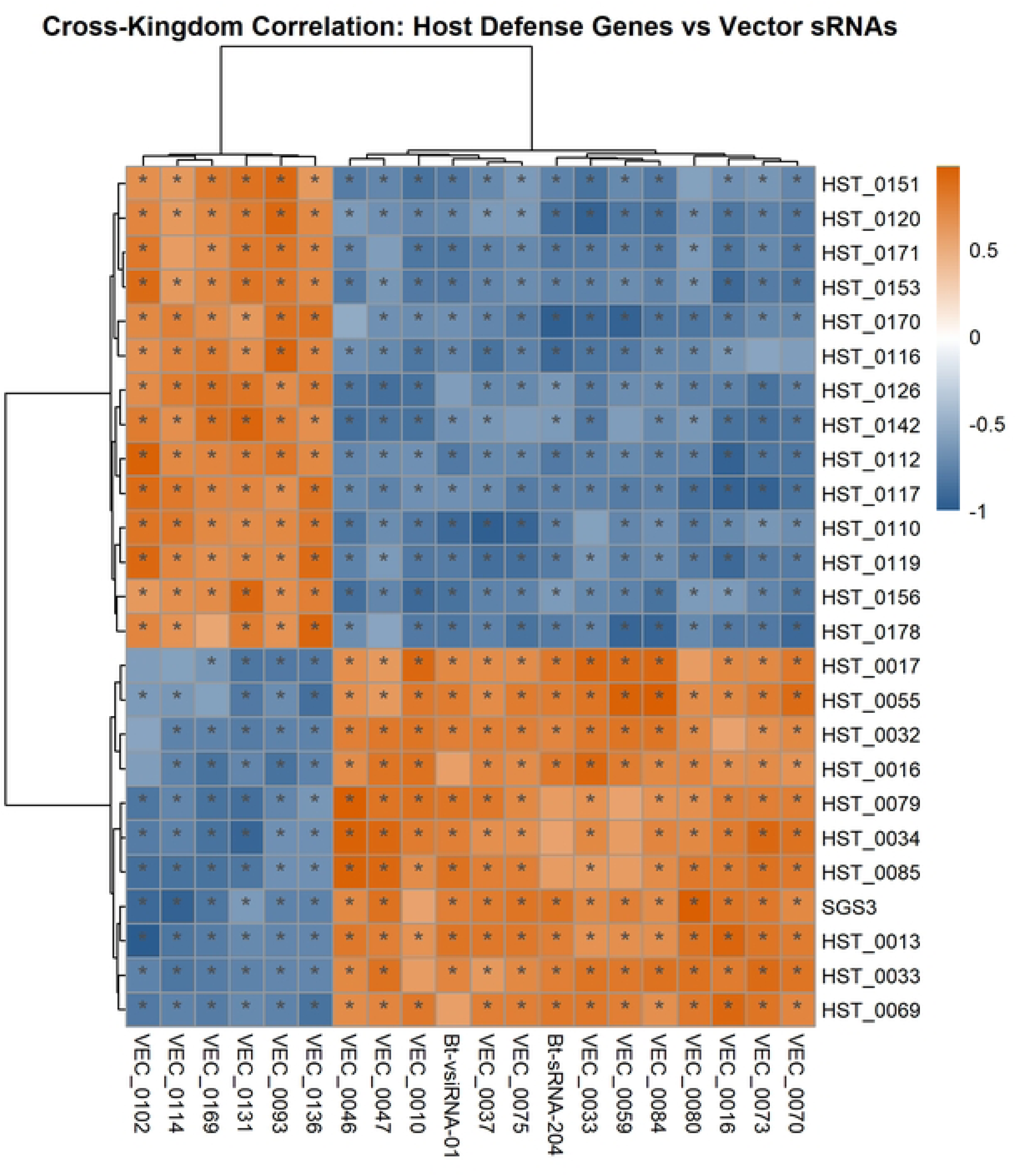
Hierarchical clustering and cross-kingdom correlation matrix between host defense features and vector small RNAs. Spearman rank correlation heatmap (ρ) mapping top 25 differentially expressed host transcripts (rows) against top 20 differentially expressed vector sRNAs and effectors (columns). Color intensity indicates direction and magnitude of co-expression, ranging from strong positive correlation (orange, ρ→ +1) to strong inverse correlation (blue, ρ→ −1). Significant cross-kingdom pairwise associations (|ρ| ≥ 0.60, P < 0.05) are designated by asterisks (*)

**Table 2.**
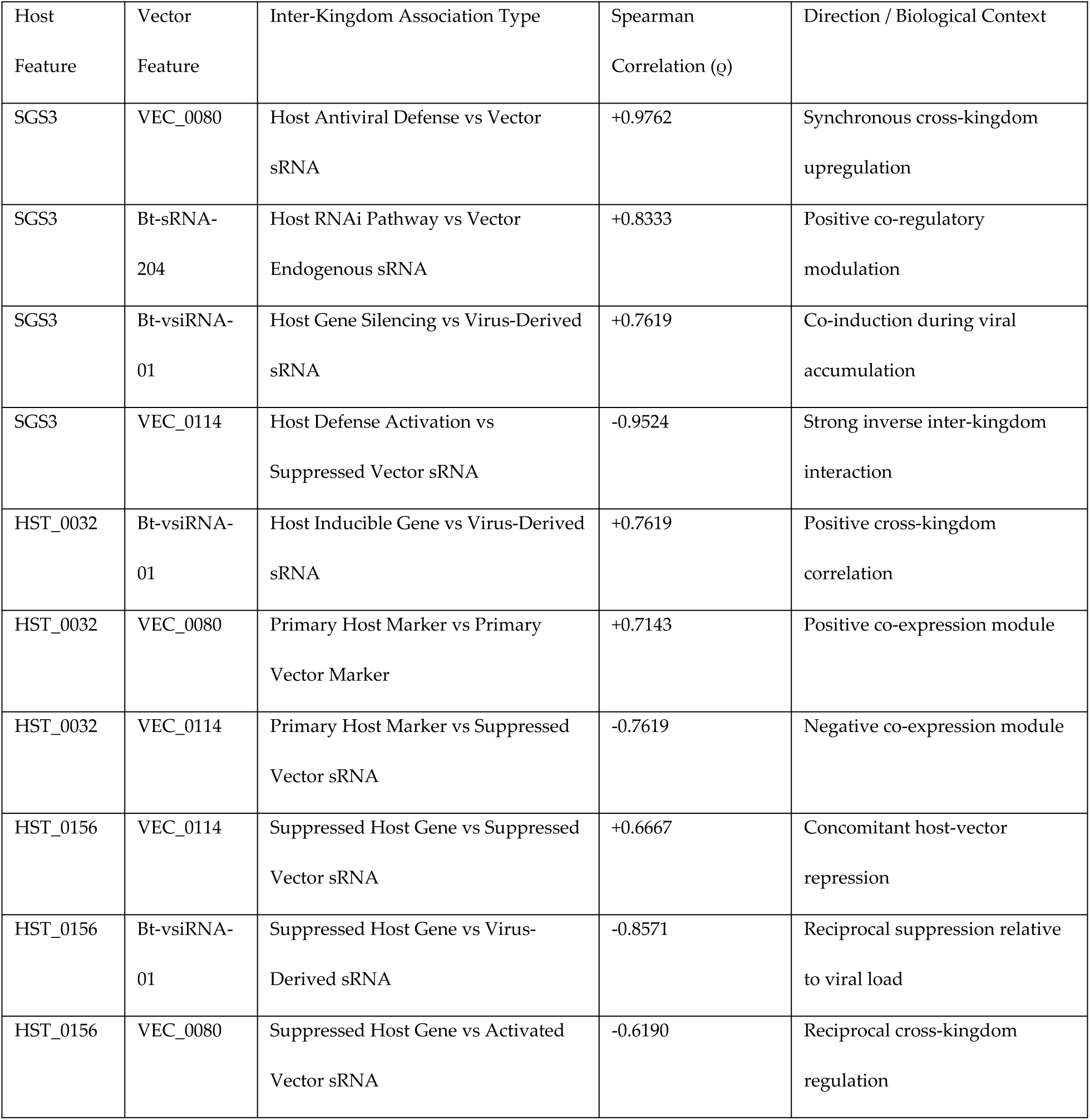
Representative cross-kingdom Spearman rank correlation pairs between host plant defense regulators and vector small RNAs.

| Host Feature | Vector Feature | Inter-Kingdom Association Type | Spearman Correlation (ρ) | Direction / Biological Context |
| --- | --- | --- | --- | --- |
| SGS3 | VEC_0080 | Host Antiviral Defense vs Vector sRNA | +0.9762 | Synchronous cross-kingdom upregulation |
| SGS3 | Bt-sRNA-204 | Host RNAi Pathway vs Vector Endogenous sRNA | +0.8333 | Positive co-regulatory modulation |
| SGS3 | Bt-vsiRNA-01 | Host Gene Silencing vs Virus-Derived sRNA | +0.7619 | Co-induction during viral accumulation |
| SGS3 | VEC_0114 | Host Defense Activation vs Suppressed Vector sRNA | -0.9524 | Strong inverse inter-kingdom interaction |
| HST_0032 | Bt-vsiRNA-01 | Host Inducible Gene vs Virus-Derived sRNA | +0.7619 | Positive cross-kingdom correlation |
| HST_0032 | VEC_0080 | Primary Host Marker vs Primary Vector Marker | +0.7143 | Positive co-expression module |
| HST_0032 | VEC_0114 | Primary Host Marker vs Suppressed Vector sRNA | -0.7619 | Negative co-expression module |
| HST_0156 | VEC_0114 | Suppressed Host Gene vs Suppressed Vector sRNA | +0.6667 | Concomitant host-vector repression |
| HST_0156 | Bt-vsiRNA-01 | Suppressed Host Gene vs Virus-Derived sRNA | -0.8571 | Reciprocal suppression relative to viral load |
| HST_0156 | VEC_0080 | Suppressed Host Gene vs Activated Vector sRNA | -0.6190 | Reciprocal cross-kingdom regulation |

The first module exhibited strong, statistically significant positive co-expression (mean *ρ* = 0.7619) between host plant antiviral defense genes (such as *SGS3*, *HST_0032*, and *HST_0033*) and viruliferous vector sRNAs/effectors (including *VEC_0080*, *Bt-vsiRNA-01*, and *Bt-sRNA-204*). For instance, host *SGS3* expression was almost perfectly correlated with vector *VEC_0080* activation and viral-derived *Bt-vsiRNA-01* accumulation (Table 2).

Conversely, the second module demonstrated strong inverse co-expression (mean *ρ* = −0.7652) between activated host defense genes and suppressed vector small RNAs (such as *VEC_0114*, *VEC_0169*, and *VEC_0093*). Furthermore, host genes downregulated during viral infection (*HST_0156* and *HST_0178*) formed a highly coherent sub-cluster that correlated positively with suppressed vector sRNAs (mean *ρ* = 0.7548) and negatively with upregulated vector sRNAs (mean *ρ* = −0.7620). These co-regulatory dynamics indicate coordinated signal transduction between plant immunity cascades and vector small RNA machinery during Begomovirus colonization.

### Pathway Enrichment Dynamics Across Host and Vector Bio-Systems

Functional enrichment analysis of differentially expressed feature sets demonstrated significant biological pathway modulation in both host plant and vector systems (Figure 4; Table 3). In host plant tissues,

**Figure 4.**
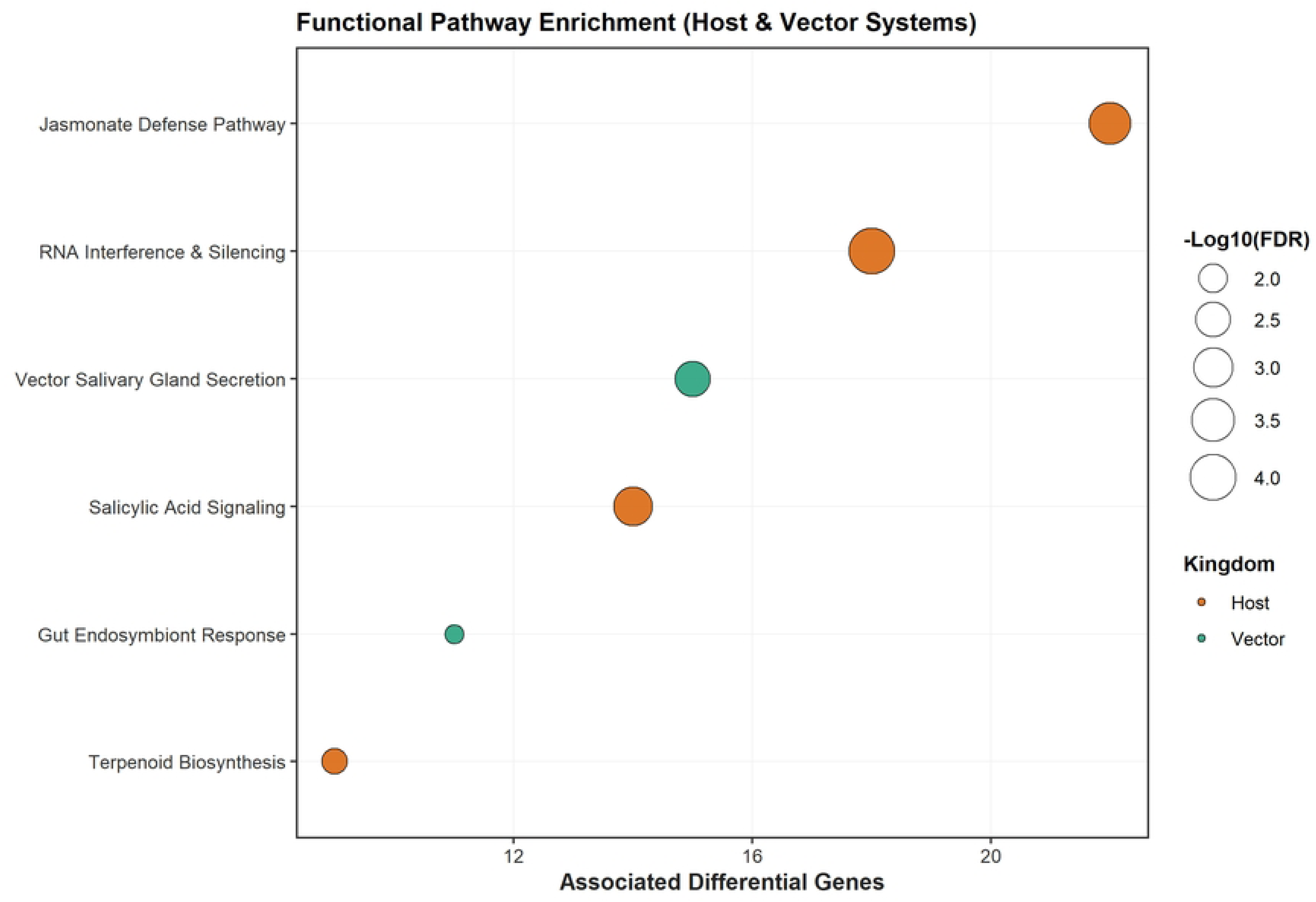
Functional pathway enrichment dynamics in host plant and insect vector bio-systems. Bubble plot displaying enriched biological pathways derived from host (orange) and vector (green) differential feature sets. The horizontal axis indicates the number of differentially expressed features associated with each pathway. Bubble size corresponds to enrichment significance expressed as -log_10_ FDR

**Table 3.**
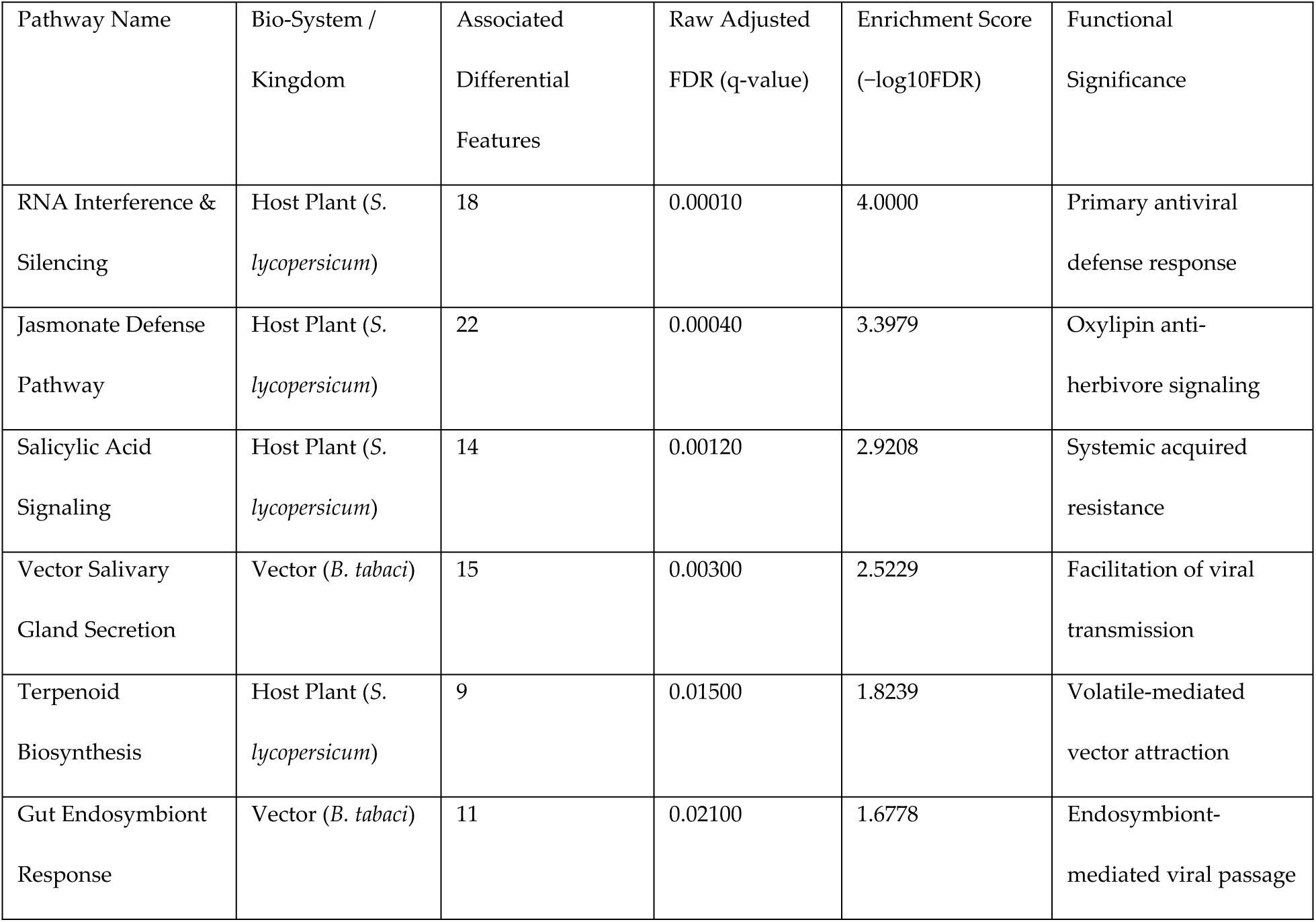
Quantitative metrics for functional pathway enrichment across host plant and vector systems.

Begomovirus infection triggered massive activation of hormone-mediated defense and gene-silencing networks:

- **Jasmonate Defense Pathway**: Highly enriched with 22 differentially expressed genes (q = 0.0004; -log_10_\FDR = 3.40).
- **RNA Interference & Silencing**: Substantially activated with 18 genes (q = 0.0001; -log_10_ FDR = 4.00), highlighting host post-transcriptional gene silencing mechanisms.
- **Salicylic Acid Signaling**: Moderately enriched with 14 genes (q = 0.0012; -log_10_ FDR = 2.92).
- **Terpenoid Biosynthesis**: Significantly altered with 9 genes (q = 0.0150; -log_10_ FDR = 1.82), reflecting secondary metabolite volatile emissions that attract vector insects.

In the insect vector (*B. tabaci*), pathway dynamics were concentrated in organs critical for viral persistence and transmission:

- **Vector Salivary Gland Secretion**: Enriched with 15 differential sRNA/transcript features (q = 0.0030; -log_10_ FDR = 2.52), pointing to alterations in salivary protein composition that facilitate viral inoculation.
- **Gut Endosymbiont Response**: Enriched with 11 features (q = 0.0210; -log_10_ FDR = 1.68), reflecting metabolic interactions between Begomovirus particles and vector primary endosymbionts (*Hamiltonella* / *Portiera*) during gut membrane passage.

## Discussion

The integration of host transcriptomic and vector sRNA datasets through a unified multi-omics pipeline successfully removed cross-platform technical variance while preserving biologically meaningful infection-driven signals. PCA demonstrated complete linear separation between infected and control samples across both kingdoms, confirming the robustness of the harmonization approach and the profound molecular impact of TYLCV infection on both host and vector. The multimodal density distributions of normalized expression intensities further validated that signal calibration achieved cross-platform comparability without introducing distributional bias. Previous studies have shown that TYLCV infection induces widespread transcriptional reprogramming in both tomato and whitefly [18,19], and our harmonization strategy enables direct cross-kingdom comparative analysis.

Differential expression analysis identified 138 significantly altered host genes, with a symmetrical distribution of 69 upregulated and 69 downregulated features. The upregulated gene set was dominated by core antiviral defense components, most notably *SGS3* (3.67-fold), which plays a critical role in the amplification of antiviral siRNAs through the RDR6-dependent pathway [20–23]. This robust induction aligns with the established function of RNAi as a primary plant defense against viruses [23]. Additional upregulated genes included *MYC2* and *PR1*, indicating activation of both JA and SA signaling pathways, consistent with the observation that begomovirus infection triggers complex hormonal defense networks [1]. The significant downregulation of photosynthetic and growth-related genes reflects a defense-growth trade-off typical of viral infections, where resources are redirected from primary metabolism to immune responses [18]. He et al. (2024) similarly reported disruption of photosynthesis-related genes and induction of immune responses in tomato during begomovirus infection [18].

In the whitefly vector, TYLCV acquisition induced substantial changes in the sRNA landscape, with 65 upregulated and 65 downregulated sRNAs. Upregulated sRNAs included both vector-encoded small RNAs and virus-derived siRNAs (Bt-vsiRNA-01), indicating that the vector’s RNAi machinery processes viral RNA intermediates during virus circulation. This is consistent with the demonstration by Shamimuzzaman et al. (2019) that virus acquisition induces piRNA clusters in whiteflies [15]. The downregulated sRNA populations suggest suppression of specific endogenous miRNAs following viral acquisition, potentially representing viral strategies to manipulate vector physiology for enhanced transmission. The pathway enrichment of vector salivary gland secretion and gut endosymbiont response factors highlights the importance of these tissues in virus transmission and persistence. Recently, researchers have reviewed how vector immunity pathways, including RNAi and other responses, serve as checkpoints for efficient begomovirus transmission [24].

The Spearman rank correlation modeling revealed two distinct, highly anticorrelated regulatory modules linking host defense genes with vector sRNAs. The correlation between host SGS3 expression and vector VEC_0080 (*ρ* = 0.9762) and Bt-vsiRNA-01 (*ρ* = 0.7619) suggests a tightly synchronized molecular dialogue across kingdoms. This near-perfect correlation implies either direct cross-kingdom signaling or shared environmental sensing mechanisms coordinating host immune activation with vector sRNA remodeling. The strong inverse correlations between activated host defense genes and suppressed vector sRNAs (mean *ρ* = −0.7652) indicate that host defenses may influence vector sRNA composition through as-yet-unidentified mechanisms. Recent study highlights that sRNAs can act across kingdoms to modulate plant–virus–vector tripartite interactions, supporting the biological plausibility of our observations [23]. The authors emphasized that non-canonical RNAi pathways and small RNAs acting across kingdoms are emerging as important modulators of these complex interactions [23].

The pathway enrichment analysis confirmed robust activation of host RNAi and JA defense pathways, with 18 and 22 differentially expressed genes respectively. The enrichment of terpenoid biosynthesis suggests that infected plants produce volatile compounds that may influence vector behavior, potentially enhancing virus spread. The enriched features in vector salivary gland secretion are consistent with the observation that begomoviruses are egested with saliva during feeding [25,26]. The gut endosymbiont response enrichment reflects the importance of whitefly endosymbionts in viral passage through the gut membrane, a critical step in circulative transmission.

Together, these results demonstrate that TYLCV infection triggers coordinated molecular responses across host and vector, with tight cross-kingdom correlation suggesting sophisticated regulatory integration. The identification of these regulatory modules provides a foundation for understanding the molecular basis of begomovirus transmission and may inform novel RNAi-based control strategies targeting both host defenses and vector transmission mechanisms. While published study support the findings of the present study, this computational analysis approach further warrants *wet lab* investigations.

## Conclusion

This multi-omics study reveals a tightly synchronized tripartite molecular crosstalk between host antiviral immunity, viral siRNA accumulation, and vector small RNA remodeling during begomovirus infection. The identification of two highly anticorrelated cross-kingdom regulatory modules, highlighted by the near-perfect correlation between host SGS3 expression and vector VEC_0080, demonstrates that host defense activation and vector sRNA responses are not independent but rather coordinated through previously unrecognized regulatory linkages. These findings advance our understanding of the complex molecular dialogue governing begomovirus transmission and suggest that vector sRNA networks are actively modulated in response to host immune status. The specific cross-kingdom regulatory modules identified here represent promising targets for dual-action RNA interference strategies aimed at simultaneously enhancing host antiviral defenses and disrupting vector transmission competence.

## Declaration Statements

## Acknowledgments

Not applicable.

## Ethics

This study did not involve human participants, human data, or human tissue. No animal experiments were conducted. All data analyzed were obtained from publicly available repositories (NCBI Gene Expression Omnibus) and did not require ethical approval.

## Data Availability

All datasets analyzed in this study are publicly available in the NCBI Gene Expression Omnibus (GEO) repository under accession numbers GSE309527 (host transcriptome profiling) and GSE111343 (vector small RNA sequencing).

## Author Contributions

A.B.: Conceptualization; Data curation; Formal analysis; Investigation; Methodology; Software; Validation; Visualization; Writing – original draft; Writing – review & editing.

K.K.: Conceptualization; Formal analysis; Investigation; Methodology; Software; Validation; Visualization; Writing – review & editing.

A.M.: Data curation; Formal analysis; Investigation; Methodology; Resources; Software; Validation; Writing – review & editing.

S.N. : Conceptualization; Project administration; Resources; Supervision; Validation; Writing – review & editing.

## References

1. Zhao SX, Wang SD, Liu YQ, Pan LL. Modulation of Plant Interactions with Whitefly and Whitefly-Borne Viruses by Salicylic Acid Signaling Pathway: A Review. Viruses. 2025;17: 825. doi:10.3390/V17060825

2. Du H, Wang YM, Wang XW. Begomovirus Transmission by the Insect Vector, the Whitefly Bemisia tabaci. Methods Mol Biol. 2025;2912: 35–47. doi:10.1007/978-1-0716-4454-6_6

3. Barboza N, Hernández E, Inoue-Nagata AK, Moriones E, Hilje L, Barboza N, et al. Achievements in the epidemiology of begomoviruses and their vector Bemisia tabaci in Costa Rica. Rev Biol Trop. 2019;67: 419–453. doi:10.15517/RBT.V67I3.33457

4. Ramos RS, Kumar L, Shabani F, Picanço MC. Risk of spread of tomato yellow leaf curl virus (TYLCV) in tomato crops under various climate change scenarios. Agric Syst. 2019;173: 524–535. doi:10.1016/J.AGSY.2019.03.020

5. Akbar A, Al Hashash H, Al-Ali E. Tomato yellow leaf curl virus (TYLCV) in Kuwait and global analysis of the population structure and evolutionary pattern of TYLCV. Virology Journal. 2024;21. doi:10.1186/S12985-024-02540-6

6. Li F, Qiao R, Yang X, Gong P, Zhou X. Occurrence, distribution, and management of tomato yellow leaf curl virus in China. Phytopathology Research. 2022;4: 28-. doi:10.1186/S42483-022-00133-1/FIGURES/3

7. Lefeuvre P, Martin DP, Harkins G, Lemey P, Gray AJA, Meredith S, et al. The Spread of Tomato Yellow Leaf Curl Virus from the Middle East to the World. PLoS Pathog. 2010;6: e1001164. doi:10.1371/JOURNAL.PPAT.1001164

8. Hasegawa DK, Shamimuzzaman M, Chen W, Simmons AM, Fei Z, Ling KS. Deep Sequencing of Small RNAs in the Whitefly Bemisia tabaci Reveals Novel MicroRNAs Potentially Associated with Begomovirus Acquisition and Transmission. Insects. 2020;11: 562. doi:10.3390/INSECTS11090562

9. Guo Q, Sun Y, Ji C, Kong Z, Liu Z, Li Y, et al. Plant resistance to tomato yellow leaf curl virus is enhanced by Bacillus amyloliquefaciens Ba13 through modulation of RNA interference. Front Microbiol. 2023;14: 1251698. doi:10.3389/FMICB.2023.1251698/FULL

10. Zhang C, Wu Z, Li Y, Wu J. Biogenesis, function, and applications of virus-derived small RNAs in plants. Front Microbiol. 2015;6: 163530. doi:10.3389/FMICB.2015.01237/XML

11. Maillard P V., Ciaudo C, Marchais A, Li Y, Jay F, Ding SW, et al. Antiviral RNA Interference in Mammalian Cells. Science. 2013;342: 10.1126/science.1241930. doi:10.1126/SCIENCE.1241930

12. Ma KW, Ma W. Phytohormone pathways as targets of pathogens to facilitate infection. Plant Molecular Biology 2016 91:6. 2016;91: 713–725. doi:10.1007/S11103-016-0452-0

13. Zhang P, Jackson E, Li X, Zhang Y. Salicylic acid and jasmonic acid in plant immunity. Hortic Res. 2025;12: uhaf082. doi:10.1093/HR/UHAF082

14. Li N, Han X, Feng D, Yuan D, Huang LJ. Signaling Crosstalk between Salicylic Acid and Ethylene/Jasmonate in Plant Defense: Do We Understand What They Are Whispering? International Journal of Molecular Sciences 2019, Vol 20, Page 671. 2019;20: 671. doi:10.3390/IJMS20030671

15. Shamimuzzaman M, Hasegawa DK, Chen W, Simmons AM, Fei Z, Ling KS. Genome-wide profiling of piRNAs in the whitefly Bemisia tabaci reveals cluster distribution and association with begomovirus transmission. PLoS One. 2019;14: e0213149. doi:10.1371/JOURNAL.PONE.0213149

16. Ritchie ME, Phipson B, Wu D, Hu Y, Law CW, Shi W, et al. Limma powers differential expression analyses for RNA-sequencing and microarray studies. Nucleic Acids Res. 2015;43: e47. doi:10.1093/nar/gkv007

17. limma: Linear Models for Microarray and RNA-Seq Data - Google Scholar. [cited 19 Oct 2023]. Available: https://scholar.google.com/scholar?hl=en&as_sdt=0%2C5&q=limma%3A+Linear+Models+for+Microarray+and+RNA-Seq+Data+&btnG=

18. He WZ, Rong T, Liu XY, Rao Q. Transcriptomic Profiling Unravels the Disruption of Photosynthesis Apparatuses and Induction of Immune Responses by a Bipartite Begomovirus in Tomato Plants. Plants. 2024;13. doi:10.3390/PLANTS13223198

19. Romero-Rodríguez B, Petek M, Kriznik M, Jiao C. Whole genome, transcriptome, smallRNAome and methylome profiling during tomato-geminivirus interaction. 2022 [cited 13 Aug 2026]. Available: https://riuma.uma.es/entities/publication/f1e300c8-2de3-4cb7-85af-1857ee353ad2

20. Tong X, Liu S, Zou J, Zhao J, Zhu F, Chai L, et al. A small peptide inhibits siRNA amplification in plants by mediating autophagic degradation of SGS3/RDR6 bodies. EMBO J. 2021;40. doi:10.15252/EMBJ.2021108050

21. Yoshikawa M, Han YW, Fujii H, Aizawa S, Nishino T, Ishikawa M. Cooperative recruitment of RDR6 by SGS3 and SDE5 during small interfering RNA amplification in Arabidopsis. Proc Natl Acad Sci U S A. 2021;118. doi:10.1073/PNAS.2102885118

22. Li F, Wang Y, Viruses XZ-, 2017 undefined. SGS3 Cooperates with RDR6 in Triggering Geminivirus-Induced Gene Silencing and in Suppressing Geminivirus Infection in Nicotiana Benthamiana. mdpi.com. [cited 13 Aug 2026]. Available: https://www.mdpi.com/1999-4915/9/9/247

23. Li F, Li X, Zhao S, Pan F, Li Z, Hao Y, et al. Antiviral RNA interference in plants: Increasing complexity and integration with other biological processes. Plant Commun. 2025;6: 101490. doi:10.1016/J.XPLC.2025.101490

24. Kuzminsky I, Ghanim M. Immunity responses as checkpoints for efficient transmission of begomoviruses by whiteflies. Virology. 2025;605: 110462. doi:10.1016/J.VIROL.2025.110462

25. Czosnek H, Hariton-Shalev A, Sobol I, Gorovits R, Ghanim M. The Incredible Journey of Begomoviruses in Their Whitefly Vector. Viruses 2017, Vol 9, Page 273. 2017;9: 273. doi:10.3390/V9100273

26. Roy B, Chakraborty P, Ghosh A. How many begomovirus copies are acquired and inoculated by its vector, whitefly (Bemisia tabaci) during feeding? PLoS One. 2021;16: e0258933. doi:10.1371/JOURNAL.PONE.0258933

